# Enhancers of host immune tolerance to bacterial infection discovered using linked computational and experimental approaches

**DOI:** 10.1101/2021.09.30.462576

**Authors:** Megan M. Sperry, Richard Novak, Vishal Keshari, Alexandre L. M. Dinis, Mark J. Cartwright, Diogo M. Camacho, Jean-François Paré, Mike Super, Michael Levin, Donald E. Ingber

## Abstract

Current therapeutic strategies against bacterial infections focus on reduction of pathogen load through antibiotics; however, stimulation of host tolerance to infection might offer an alternative approach. Here we used computational transcriptomics and a *Xenopus* embryo infection model to rapidly discover infection response pathways, identify potential tolerance inducer drugs, and validate their ability to induce broad tolerance. *Xenopus* embryos exhibit natural tolerance to *A. baumanii, K. pneumoniae, S. aureus*, and *S. pneumoniae* bacteria, whereas *A. hydrophila* and *P. aeruginosa* produce infection that leads to death. Transcriptional profiling led to definition of a 20-gene signature that allows for discrimination between tolerant and susceptible states, as well as identification of active and passive tolerance responses based on the degree of engagement of gene transcription modulation. Upregulation of metal ion transport and hypoxia pathways reminiscent of responses observed in primate and mouse infection models were identified as tolerance mediators, and drug screening in the susceptible *A. hydrophila* infection model confirmed that a metal chelator (deferoxamine) and HIF-1α agonist (1,4-DPCA) increase embryo survival despite high pathogen load. These data demonstrate the value of combining the *Xenopus* embryo infection model with multi-omics analyses for mechanistic discovery and drug repurposing to induce host tolerance to bacterial infection.

## 1. Introduction

Current therapeutic strategies against bacterial infection focus on direct killing of bacteria with antibiotics or induction of immunological resistance through use of vaccines, each of which works by reducing pathogen load.^[1,2]^ These strategies have proven highly effective in the past, but they also have substantial disadvantages and face major challenges as development of bacterial resistance to antibiotics is becoming rampant and people must be willing to be vaccinated prior to infection to confer any defense against disease. An alternative approach is to develop therapeutics that act by augmenting host tolerance to infection in the continued presence of pathogens, rather than killing or inactivating the microbes.^[1,2]^ This possibility is supported by the observation that different species exhibit a wide range of tolerance to infection with the same pathogen.^[3–5]^ For example, recent work suggests that mice achieve tolerance in their nasopharynx relative to other anatomical sites during pneumococcal infection.^[6]^ Tolerance is also seen in higher order mammals with African and Asian monkeys (AAM) exhibiting greater natural tolerance compared to humans and baboons when exposed to immune stimulation with bacterial lipopolysaccharide endotoxin (LPS).^[7]^ Thus, it may be possible to develop therapeutics that similarly induce a state of host tolerance to minimize infection-related organ injury and systemic disease, which could be used as a preventative measure when entering a highly infection-prone environment or even to prolong the survival of the patient until proper treatment is available.

The mechanisms underlying disease tolerance seem to revolve around a sophisticated network of evolutionarily conserved stress responses that initiate tissue damage control in the infected host.^[8]^ Initiation of stress responses depends on the activation of sensors that monitor environmental conditions and homeostasis in the host, including oxygen levels, pH, glucose, and ATP.^[1,9]^ Experimental models suggest that host tolerance is associated with cellular hypoxia and upregulation of hypoxia inducible factors (HIFs), which reprogram leukocyte metabolism and control the balance of TH17/Treg cells, thereby limiting T cell-induced pathology.^[10]^ Oxygen levels also regulate iron uptake and copper efflux,^[8]^ which in turn regulate macrophage function.^[11]^ In addition, metabolic reprogramming in host cells and tissues deprives infectious pathogens of essential nutrients, such as transition metals required for the function of metalloproteinases and other enzymes. Because tight regulation of iron, zinc, manganese, and copper uptake is required for pathogenic bacteria to thrive, shifts in host metabolism can produce metal starvation by sequestration or induce bacterial toxicity by releasing high concentrations of metals.^[12]^

Despite increasing knowledge about the complex network of mechanisms driving tolerance in experimental models of infection, the development of therapeutics to target these mechanisms has been limited. We and others have demonstrated the usefulness of drug repurposing approaches using *in vitro* models, such as human organs-on-chips, to rapidly identify drug candidates that can be advanced into further pre-clinical testing and human clinical trials.^[13–15]^ In this study, we set out to explore if we can use integrated and iterative bioinformatics and experimental approaches, including leveraging *Xenopus* embryos that have not yet developed an adaptive immune system, to discover immune tolerance induction pathways, repurpose existing FDA approved drugs, and carry out efficacy and toxicity screening assays to identify compounds that induce tolerance to bacterial infection. Importantly, we evaluated host tolerance in the context of six clinically relevant bacterial infections to maximize translatability and performed infection and drug screening studies in *Xenopus* frog embryos, which have close homology with the human genome.^[16]^ In infection-tolerant embryos, we identified the activation of genes involved in metal ion binding, membrane transport, and oxygen transport pathways. Based on these findings, we repurposed existing metal ion scavengers and a HIF agonist that artificially induce tolerance to infection, prevent transcriptional changes associated with infection, and increase *Xenopus* embryo survival.

## 2. Results

### 2.1 Xenopus embryos demonstrate natural tolerance to multiple bacterial infections

Similar to humans and other organisms,^[3]^ *Xenopus* frog embryos demonstrate natural tolerance to high bacterial loads of some microorganisms.^[17]^ We confirmed this by testing the *Xenopus* response to infection with six clinically-isolated bacterial pathogens (*A. baumanii, K. pneumoniae, S. aureus, S. pneumoniae, A. hydrophila*, and *P. aeruginosa*) using topical application or direct microinjection and found that the embryos tolerate infections caused by *A. baumanii, K. pneumoniae, S. aureus*, and *S. pneumoniae* bacteria without outward adverse effects (**Figure 1** & **Figure S1**). More than 80% of embryos survived infection with these microorganisms for up to 52 hours (**Figure 1A**), while carrying median bacterial loads of 4.1-8.5 × 10^4^ CFU (**Figure 1B**). In contrast, infection with *A. hydrophila* and *P. aeruginosa* at a similar pathogen burden led to death of *Xenopus* embryos (**Figure 1A**), suggesting that embryos exposed to these bacterial species are unable to shift to a tolerant state. The outward appearance of infection-tolerant embryos was indistinguishable from uninfected embryos, whereas development was arrested in susceptible embryos, which did not survive longer than 52 hours post-infection (**Figure 1C**).

**Figure 1.**
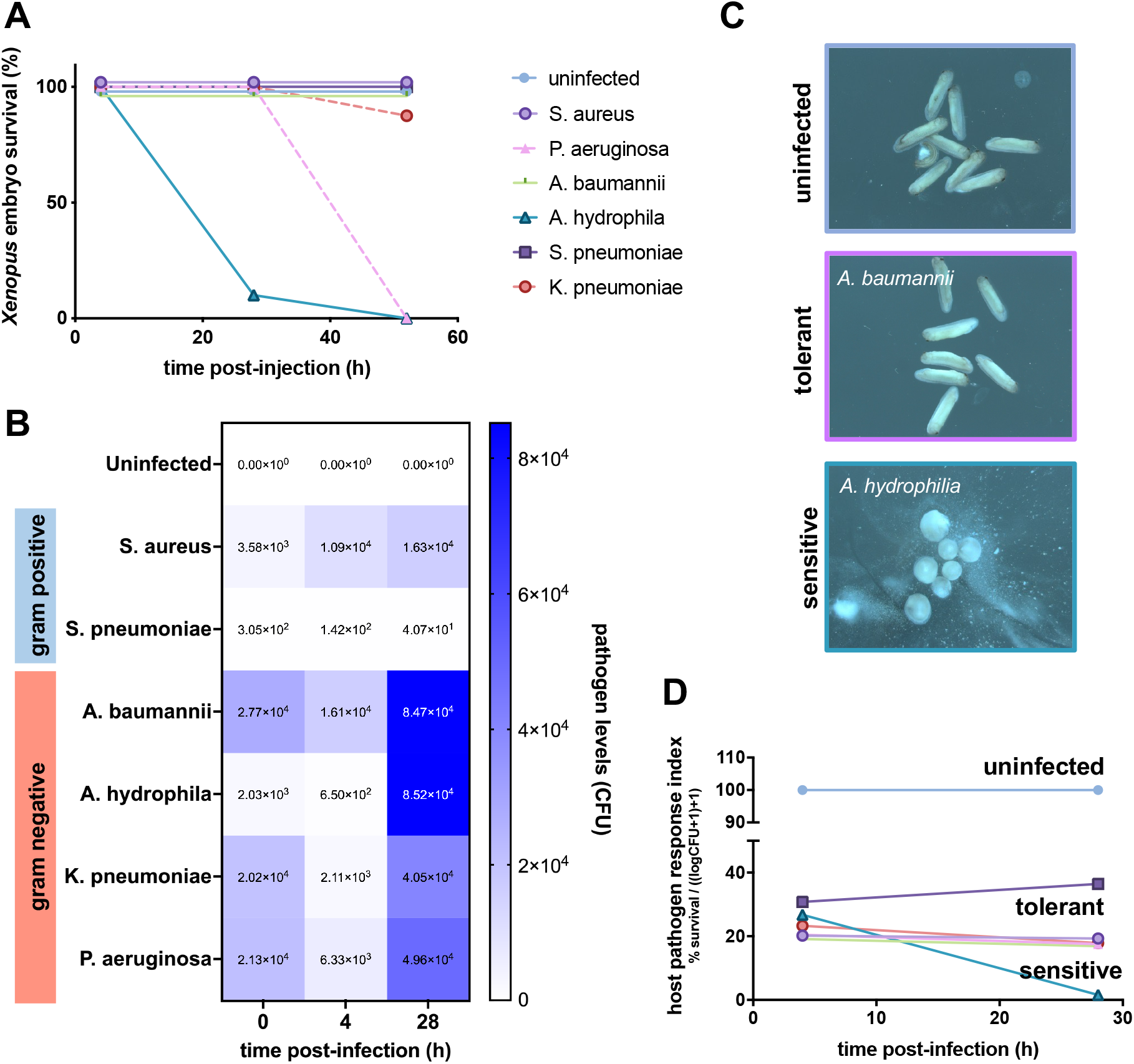
*Xenopus* embryos demonstrate natural tolerance to high bacterial loads of some pathogens. **(A)** Embryos survive after microinjection of high bacterial loads of *A. baumanii, K. pneumoniae, S. aureus*, and *S. pneumoniae* without outward adverse effects. **(B)** Pathogen levels are elevated over control levels in both tolerant and sensitive embryos. **(C)** At 52 hours post-infection, the morphology of infection-tolerant embryos (*A. baumannii*-infected shown) is indistinguishable from uninfected embryos, whereas development is arrested in susceptible embryos (*A. hydrophila*-infected shown). **(D)** The host pathogen response index combines survival and bacterial load data, thus permitting differentiation of resistant, tolerant, and susceptible states in *Xenopus* embryos. (n=10 embryos/group)

Although the evaluation of morphology and survival metrics suggest that infection-tolerant embryos are ostensibly similar to their uninfected counterparts, we found that evaluation of both survival and infection burden provided the clearest measurement of an embryo’s biological state, as previously observed by others.^[17]^ To better quantify the host’s infection tolerance state, we therefore developed a host-pathogen response index (HPRI) that captures both the host response and the pathogen load in a single score by normalizing the percent embryo survival by the pathogen burden (**Figure 1D**). Calculation of the HPRI metric revealed that infection-tolerant embryos exhibit a host-pathogen response trajectory distinct from both the uninfected and susceptible groups. Additionally, the differences in HPRI between *S. pneumoniae* (HPRI=36) and other host-tolerant infections (HPRI=17-19) as early as 24 hours post-infection suggests that multiple states of host tolerance might exist, similar to the classes of tolerance outlined by Ayres & Schneider (2012).^[2]^ Since embryos infected with *S. pneumoniae* have a lower pathogen load compared to other host-tolerant infections, it is also possible that *Xenopus* exhibit some degree of resistance to *S. pneumoniae*.

### 2.2 Gene expression analysis stratifies active and passive tolerance states

To define the gene circuits and pathways involved in the infection-tolerant state, we measured the gene expression of embryos infected with each of the six types of bacteria both early after exposure (4 hours) and 1 day later (28 hours) when the first impacts of infection on embryo health are observed (**Figure 1**). At 4 hours, minimal changes in gene expression were observed compared to the uninfected control state (**Figure 2A**). As expected from the survival curves, by 28 hours post-infection we detected widespread changes in gene expression in embryos exposed to *A. hydrophila* and *P. aeruginosa* that are sensitive to infection. Notably, hierarchical clustering of gene expression stratified the infection-tolerant groups into an “active” tolerance group that induced a large shift in gene expression (*A. baumanii* and *K. pneumoniae*) and a “passive” tolerance group that produced more subtle gene changes (*S. aureus* and *S. pneumoniae*) similar to those observed at the 4-hour time point (**Figure 2A**).

**Figure 2.**
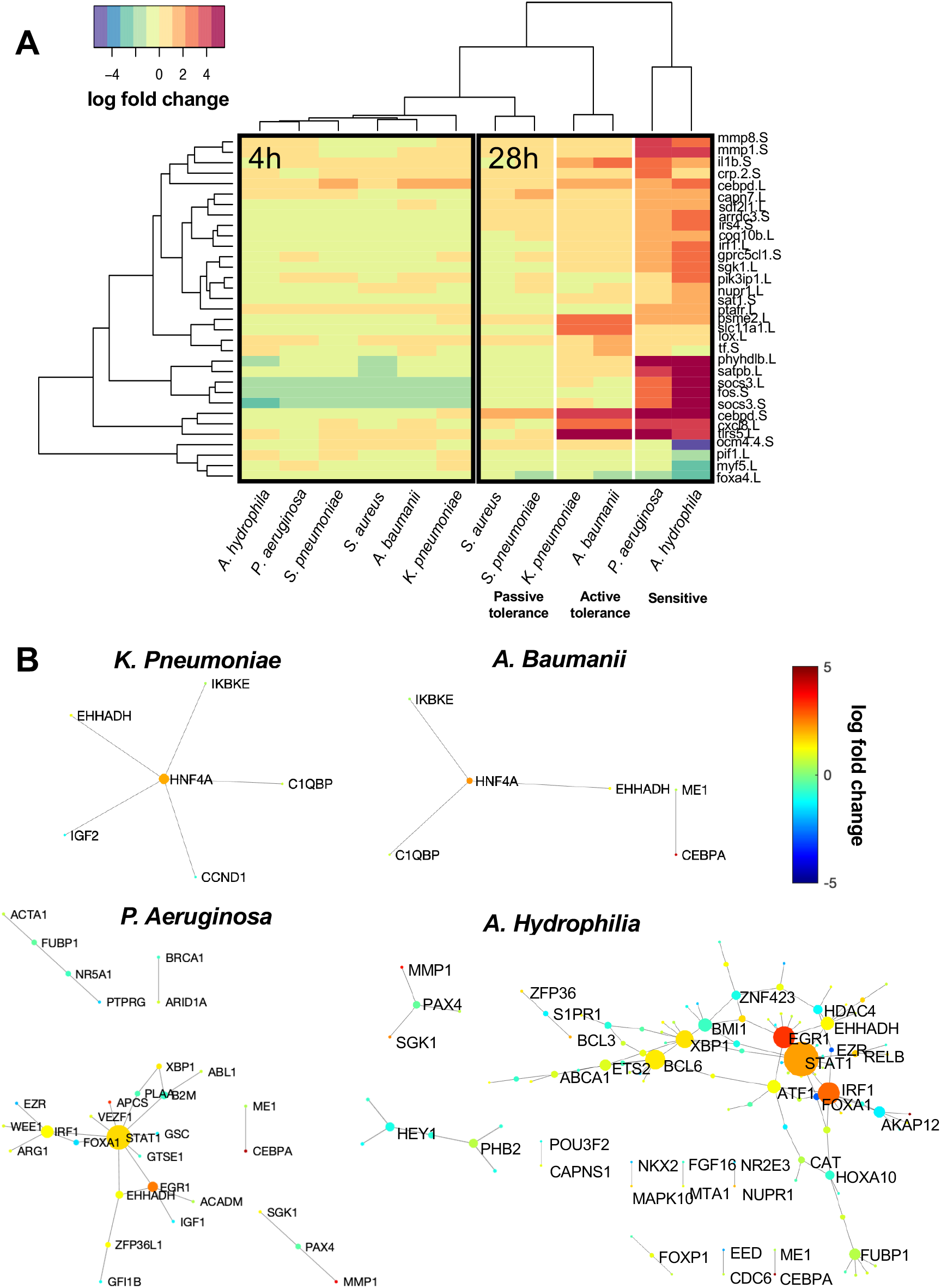
Gene expression analysis stratifies active and passive tolerance states. **(A)** Heatmap comparing the expression of 33 genes that undergo significant changes in gene expression relative to uninfected controls at matching time points (padj<10^−4^). Hierarchical clustering analysis reveals that infection begins as a generic infection response at 4 h post-infection and at 28 h proceeds into stratification by tolerance and into active and passive tolerance states, with *A. baumanii* and *K. pneumoniae* stimulating substantial gene upregulation with fold change greater than 2 (active tolerance) and *S. aureus* and *S. pneumoniae* infection leading to only minor changes in gene expression (passive tolerance). **(B)** Active tolerance and susceptible states are characterized by the activation of gene-gene subnetworks, whereas no subnetwork activation was observed in the passive tolerance states. Genes within these networks undergo significant changes in gene expression (padj<0.05) and edges are defined by known interactions as defined by the TRRUST and KEGG databases. Gene node size corresponds to that gene’s degree of interconnectedness with other genes in the network. Gene node color designates the fold change in gene expression.

To understand if and how gene subnetworks and circuits may be reorganized in the tolerant state compared to sensitive, we mapped *Xenopus* genes to their human orthologs and identified interconnected networks of significantly changed genes based on known interactions from the Kyoto Encyclopedia of Genes and Genomes (KEGG) and Transcriptional Regulatory Relationships Unraveled by Sentence-based Text mining (TRRUST) databases.^[18–20]^ Inspection of these interconnected networks of differentially expressed genes revealed a shift in gene sub-networks in the sensitive compared to active tolerance states with the infection tolerance state being characterized by the upregulation of HNF4A in embryos exposed to *A. baumanii* and *K. pneumoniae*, and this gene was also identified as a major hub of the active tolerance gene network (**Figure 2B**). A common motif comprised of the genes IKBKE, C1QBP, and EHHADH was found to be directly connected to the HNF4A hub in the infection tolerant state associated with *A. baumanii* and *K. pneumoniae* infection. IKBKE and C1QBP are known to regulate inflammatory and infection processes, with hematopoietic IKBKE limiting inflammasome priming and metaflammation.^[21]^ EHHADH regulates fatty acid oxidation, a metabolic pathway with reduced activity during infection due to remodeling of molecular components, impacting innate immunity and host tolerance.^[22,23]^ In addition, the primary hub gene HNF4A represses CLOCK-ARNTL/BMAL1 transcriptional activity and is essential for circadian rhythm maintenance and period regulation in liver and colon cells,^[24]^ which is reminiscent of the importance of endogenous rhythms for survival during influenza infection.^[25]^

In contrast, the sensitive state was characterized by much more expansive gene sub-networks, with the transcription factors STAT1, EGR1, and IRF1 being upregulated and acting as highly connected transcriptional network hubs in embryos exposed to *A. hydrophila* and *P. aeruginosa* (**Figure 2B**). No sub-networks were identified for embryos treated with *S. aureus* and *S. pneumoniae*, suggesting that passive tolerance is minimally different from the uninfected state or, in other words, that these bacteria fail to interact biologically with the host in a significant way (**Figure 1D**), rather than inducing a tolerance gene program.

### 2.3 An active tolerance gene signature associated with metal ion binding and transport

Our transcriptional analyses allowed us to identify a novel signature for infection tolerance, comprised of 20 unique genes. Briefly, we compared the expression profiles between controls and the tolerant or sensitive samples. This allowed us to define a tolerance signature defined by the set of genes that were differentially expressed in the tolerant cases but did not change in the sensitive cases (**Figure 3A**). We then mapped the tolerant-specific *Xenopus* genes to human orthologs and used PANTHER^[26]^ to perform functional classification analysis of genes using gene ontology (GO) and Reactome pathway databases.^[27–29]^ GO analysis showed that these genes are involved in pathways related to ion binding, particularly the binding of cyclic compounds, cations, transition metal ions, and zinc (**Figure 3B**). Evaluation of Reactome pathways similarly identified pathways involved in small molecule transport, G(q) signaling, and solute carrier (SLC)-mediated transport. In addition to genes with ion binding and membrane transport functions, the 20-gene tolerance signature also includes the gene NGB, which is upregulated in infection-tolerant embryos and is involved in increasing oxygen availability, which provides protection under hypoxic and ischemic conditions.^[30]^ The gene HNF4A, which was identified as a major gene network hub in tolerance networks, is also the most upregulated component of the gene signature.

**Figure 3.**
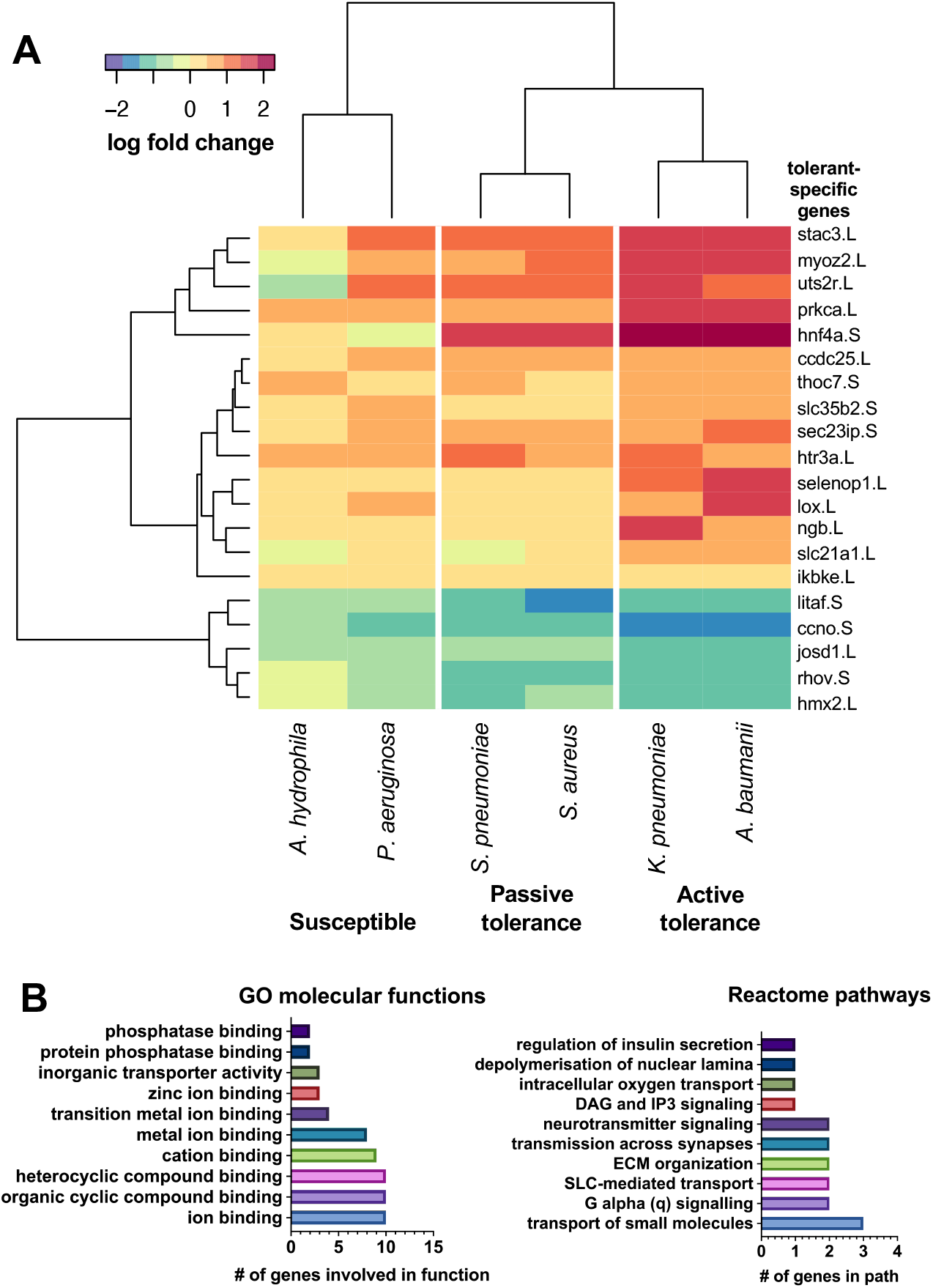
Genes involved in metal ion binding, transport and signaling, and ECM organization define the active tolerance state. **(A)** A heatmap of the 20-gene signature that defines the host tolerance state in *Xenopus* embryos. These genes are up or down regulated in the tolerance states (padj<0.05) and are unchanged in the susceptible state. **(B)** Genes within the tolerant signature are involved in multiple gene ontology (GO) and Reactome pathways. Ion binding pathways, including metal, cyclic compound, and cations, are strongly represented within the tolerance signature. In addition, biological transport and signaling pathways are prominently represented, including intracellular oxygen transport.

### 2.4 Gene network reorganization is detected in both Xenopus and mammalian infection

To assess if our findings in the *Xenopus* infection model are more broadly generalizable to mammals, we compared *Xenopus* gene expression during infection tolerance to that in mice and primates from previously published work. First, we identified genes that are differentially expressed across tolerance states in *Xenopus*, the nasopharynx of mice infected with *S. pneurmoniae*,^[6]^ and primates exposed to LPS^[7]^ and then we evaluated differentially expressed gene networks to identify global activity patterns and common motifs. Comparison of infection tolerance transcriptomic signatures in mice and primates relative to *Xenopus* reveals overlapping features, as well as unique aspects of tolerance in different species and tissues (**Figure 4**). We identified 12 genes that are differentially expressed in at least one tolerant condition for each of the species studied (**Figure 4A**), amongst which there are multiple genes involved in nuclear factor kappa B (NF-κB) signaling, metal ion transport, and cellular stress responses. Similar to *Xenopus*, tolerance in the mouse nasopharynx and primates exposed to LPS is characterized by upregulation of the genes IKBKE, NFKBIA, and BCL3, which are involved in the NF-κB signaling pathway. Both IKBKE and NFKBIA encode for inhibitors of NF-κB and are involved in regulating inflammatory responses to infection. IKBKE specifically plays an important role in energy balance regulation, including limiting chronic inflammation during metabolic disease and atherosclerosis.^[21]^ BCL3 is a pro-survival and antiapoptotic gene.^[31]^ LCP2, which encodes for lymphocyte cytosolic protein 2, is also upregulated across *Xenopus* and mammals studied.

**Figure 4.**
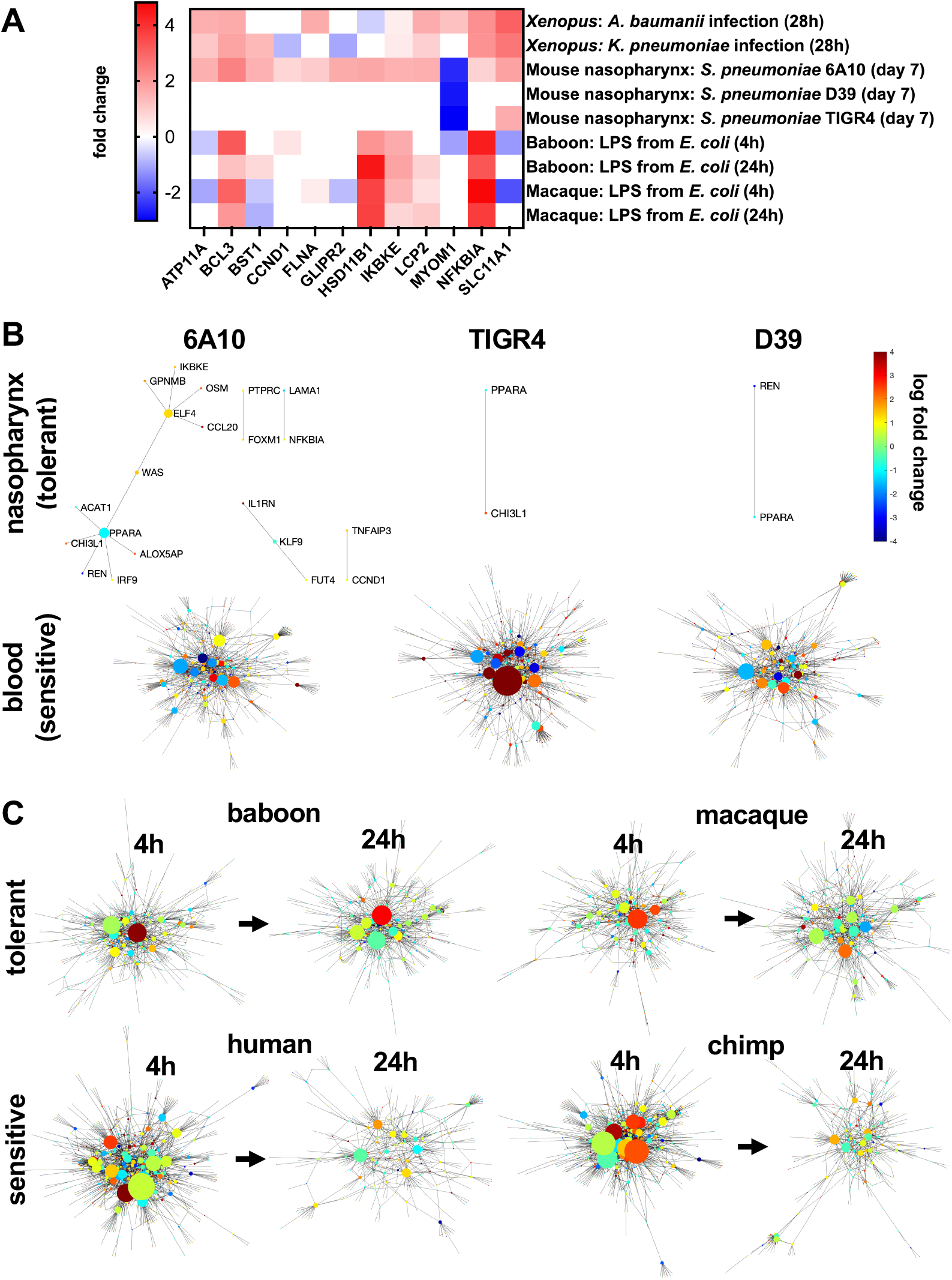
Cross-species analysis of infection tolerance. **(A)** Differentially expressed genes (padj<0.05) common to *Xenopus*, mouse, and primate tolerance states. Shaded squares represent significantly differentially expressed genes and blank squares indicate that the gene is not differentially expressed for a given condition. Gene-gene networks comprised of differentially expressed genes for tolerant and sensitive states in **(B)** mice exposed to *S. pneumoniae* and **(C)** primates exposed to LPS from *E. coli*. In mice, the nasopharynx is an anatomical site of tolerance compared to the more sensitive environment in the blood in response to *S. pneumoniae*. Baboons and macaques exposed to LPS exhibit greater tolerance compared to humans and chimps, especially at the 4h time point.

Genes involved in metal ion transport, including FLNA, SLC11A1, ATP11A, and CCND1 are also differentially expressed in tolerance states across species (**Figure 4A**). FLNA is upregulated in all species assessed, however, SLC11A1 and CCND1 exhibit different expression levels across species. SLC11A1, a divalent transition metal (iron and manganese) transporter, and ATP11A, a membrane ATPase, are upregulated in tolerant *Xenopus* and mice but are downregulated in primate disease tolerance. CCND1, a known core effector gene involved in repressing transcriptional stress and damage responses,^[32]^ is upregulated in mammals and is downregulated in *Xenopus*, possibly due to differences in tissue types and species-specific responses.

In the mouse nasopharynx, gene networks of differentially expressed genes are less widespread compared to networks observed in blood samples (**Figure 4B**). Notably, the genes IKBKE and CCND1 are involved in mouse as well as *Xenopus* tolerance gene networks. However, these *Xenopus* and mouse nasopharynx networks exhibit distinct features as well, such as PPARA, which acts as a hub node in tolerance networks of the mouse nasopharynx across pathogen strains and is not implicated in *Xenopus* tolerance networks. Interestingly, PPARA deficiency in mice is lethal in cases of infection with LPS-producing bacteria.^[23]^

In contrast to the smaller gene networks activated in tolerant *Xenopus and* mouse, primates exposed to LPS exhibit much more widespread network activation (**Figure 4C**). Activation of gene networks occurs rapidly in both tolerant and sensitive primates, following only 4h of exposure to LPS. At this same time point, infected *Xenopus* exhibited only subtle differences in gene expression and no gene subnetwork activation (**Figure 2**). Widespread differential gene networks are also present in primates after 24h of LPS exposure. In tolerant primates, a greater number of strong hub genes are observed in gene networks at 24h compared to those for sensitive primates, suggesting high interconnectivity in tolerance networks (**Figure 4C**). Assessment of differentially expressed genes that are shared among tolerant primates but not among sensitive primates revealed network modules comprised of genes involved in interferon alpha (5/198 genes) and gamma (7/198 genes) signaling, matching a past analysis.^[7]^ Substantial portions of the gene network are involved in T cell activities, including activation (11/198 genes), migration (6/198 genes), and differentiation (10/198 genes) (**Figure S2**), which were not widely identified in *Xenopus* analyses of tolerance likely due to the relative immaturity of their immune system.^[17]^ However, similar to the *Xenopus* response, a substantial portion of genes in the primate tolerance network are involved in responses to metal ions (12/198 genes), oxygen (8/198 genes), and hypoxia (12/198 genes). Together, the overlaps in active gene subnetworks with mice and primates support the possibility that many of the findings we obtained in the *Xenopus* infection model may translate to mammalian models and humans.

### 2.5 Metal ion scavenging and hypoxia-promoting drugs improve infection tolerance

Based on our characterization of the active tolerance state (**Figures 3 and 4**), we screened drugs that modulate metal ion transport or hypoxia (key compounds outlined in **Table 2**) in the susceptible *A. hydrophila* infection model to explore if they could shift the embryos to a state of infection tolerance (**Figure S1**). First, we tested the iron and aluminum ion chelator drug deferoxamine (DFOA), which is approved by the FDA for treatment of hemochromatosis. Importantly, DFOA increased embryo survival over 120 hours of infection in a dose-dependent manner (**Figure 5A**). While there was a small decrease in pathogen load with the highest concentration of DFOA (2 mM), the lower 0.2 mM dose was able to significantly increase survival in the presence of a sustained high pathogen load that was similar to that seen in the infected controls. Similar results were achieved with the iron and zinc scavenger, L-mimosine (**Figure S3**), which substantially increased embryo survival, but minimally impacted pathogen load. The vasodilator hydralazine, which also complexes with metal ions, likewise improved embryo survival, but over a smaller range of concentrations (**Figure S3**).

**Figure 5.**
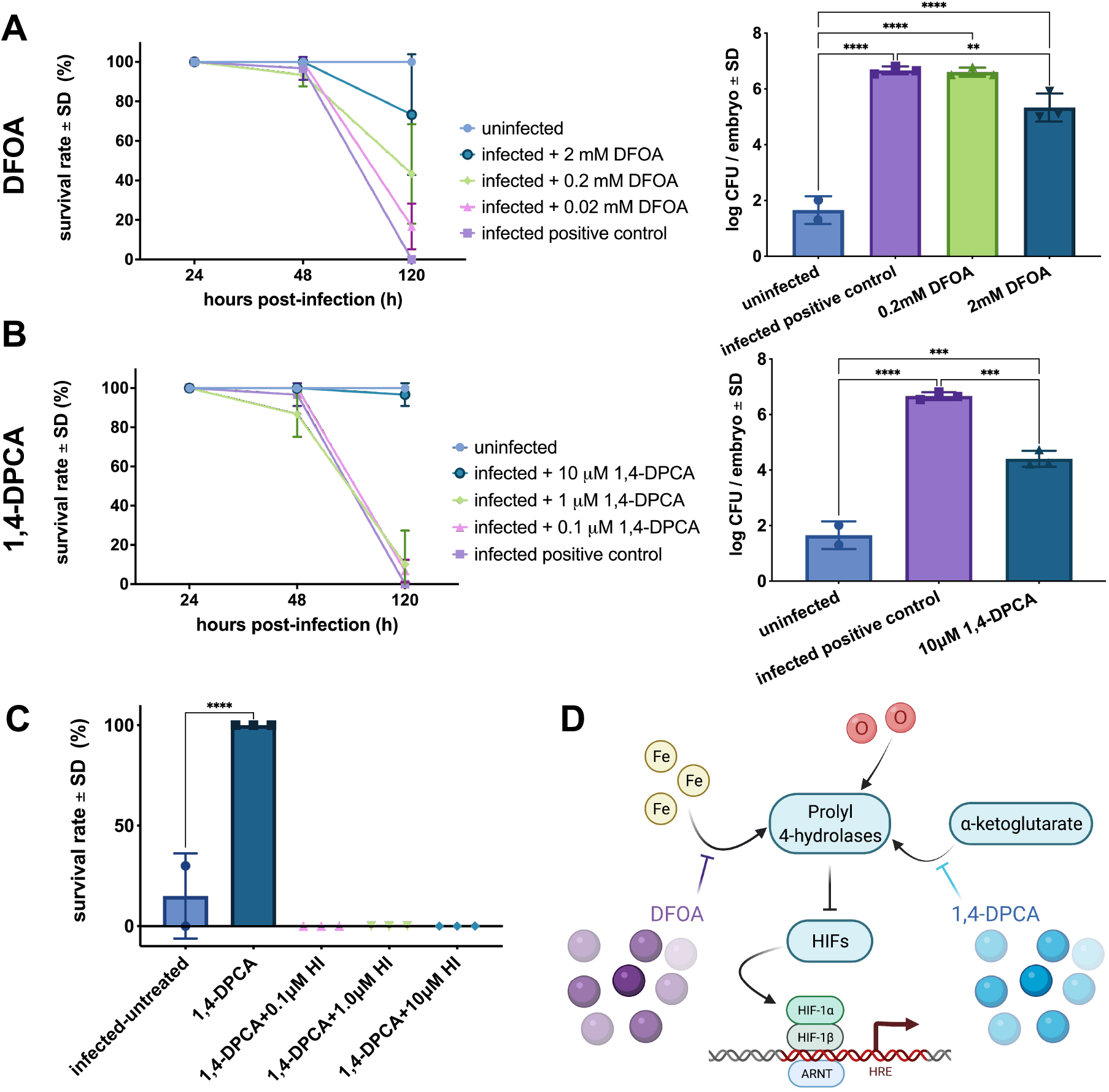
Metal ion transport and hypoxia-modulating drugs improve infection tolerance. **(A)** The iron and aluminum scavenger deferoxamine (DFOA) improves survival in *Xenopus* embryos in a dose-dependent fashion, with greater than 75% survival after 2mM DFOA treatment (n=3 replicates/group/time point with n=10 embryos each). 2mM DFOA also reduces pathogen burden; however, CFU levels remain higher than uninfected embryos at 120 hours post-infection (**p=0.009; ****p<0.0001). Statistical analysis was performed using a one-way ANOVA of the log-transformed data and a Tukey’s multiple comparisons test. **(B)** The HIF-1α agonist 1,4-DPCA also increases embryo survival when dosed at 10μM (n=3 replicates/group/time point with n=10 embryos each). Similar to DFOA treatment, 10μM 1,4-DPCA reduces pathogen load, but not to uninfected levels (***p<0.001; ****p<0.0001). Statistical analysis was performed using a one-way ANOVA of the log-transformed data and a Tukey’s multiple comparisons test. **(C)** Embryos are reversed to a susceptible state by HIF-1α inhibition during treatment with 1,4-DPCA and no survival is observed in the inhibition groups (****p<0.0001; n=3 replicates/group/time point with n=10 embryos each). Statistical analysis was performed using a one-way ANOVA and post-hoc comparison of each condition to the infected-untreated group. **(D)** Pathway model linking HIF and metal ion transport mechanisms.

We hypothesized that metal ion binding may be inducing a hypoxia response via HIF-1α. When we tested the HIF-1α agonist, 1,4-DPCA, which inhibits prolyl-4 hydroxylase by competing with its α-ketoglutarate substrate, we found that at a dose of 10μM it increased embryo survival while maintaining a high pathogen load (**Figure 5B**). Similar to the effects of the highest dose of DFOA, pathogen load decreased slightly after 1,4-DPCA treatment relative to the infected control embryos, but not to uninfected levels (**Figure 5A**,**B**). To confirm the role of hypoxia pathways in infection tolerance, we explored whether we could shift infected embryos to a susceptible state during 1,4-DPCA treatment using the research-grade HIF-1α inhibitor, sc-205346. Indeed, inhibition of HIF-1α activity reversed the embryo rescue observed during 1,4-DPCA treatment alone, even when the HIF-1α inhibitors were dosed at low concentrations (**Figure 5C**). Together, these compound screens suggest that infection tolerance may be achieved by targeting proly-4 hydroxylase to induce the hypoxia pathway (**Figure 5D**).^[8]^

Based on the upregulation of circadian transcription factor HNF4A and its role as a gene network hub in the tolerance state (**Figures 2 & 3)** as well as prior work linking hypoxia to circadian rhythm,^[33]^ we also explored the link between circadian rhythm and tolerance. We first tested if 1,4-DPCA is capable of inducing tolerance across the circadian cycle and observed that the drug worked equally well regardless of light cycle (**Figure S4A**). Unexpectedly, the flipping of light cycle *alone* induced *Xenopus* tolerance to *A. hydrophila* infection (**Figure S4B**). This unanticipated finding suggests that infection tolerance is controlled by multiple intersecting pathways, including the three explored in this study: hypoxia, metal ion transport, and circadian rhythm.

### 2.6 1,4-DPCA treatment mimics active infection tolerance

Given our finding that drug exposure impacts both the host and pathogen, we set out to distinguish the effects of 1,4-DPCA on the pathogen versus the host. Using susceptible embryos infected with *A. hydrophila*, we assessed the survival and pathogen burden of embryos treated prophylactically or after infection (**Figure 6** & **Figure S1**). In the prophylactic treatment groups, we explored the effects of pretreating the pathogen or the *Xenopus* embryos separately to assess differential role of the drug on the overall tolerance response. Prophylactic treatment of either the pathogenic microbes or both the embryo and the pathogens simultaneously resulted in an increase in embryo survival to levels similar to that of uninfected controls (**Figure 6A**). Prophylactic treatment of embryos alone was less effective, but survival levels improved relative to untreated infection. Pathogen load decreased across all prophylactic groups except in the case of prophylactic treatment of embryos alone. However, bacterial CFUs were detectable in all prophylactically treated embryos whereas they were not in uninfected controls. The HPRI score, which combines these metrics, revealed that regardless of the prophylactic administration regimen utilized, all treated embryos improve over untreated infection but maintain a lower HPRI relative to uninfected controls. This result suggests that prophylaxis with 1,4-DPCA induces some level of tolerance, but not resistance, in embryos infected with *A. hydrophila*. Importantly, treatment after the initiation of infection also substantially improved embryo survival with > 80% of embryos surviving infection (**Figure 6B**). Pathogen load decreased across all post-infection treatment embryos and the HPRI score also improved for these groups, with the best results observed in embryos exposed to a combination of prophylactic and post-infection treatments.

**Figure 6.**
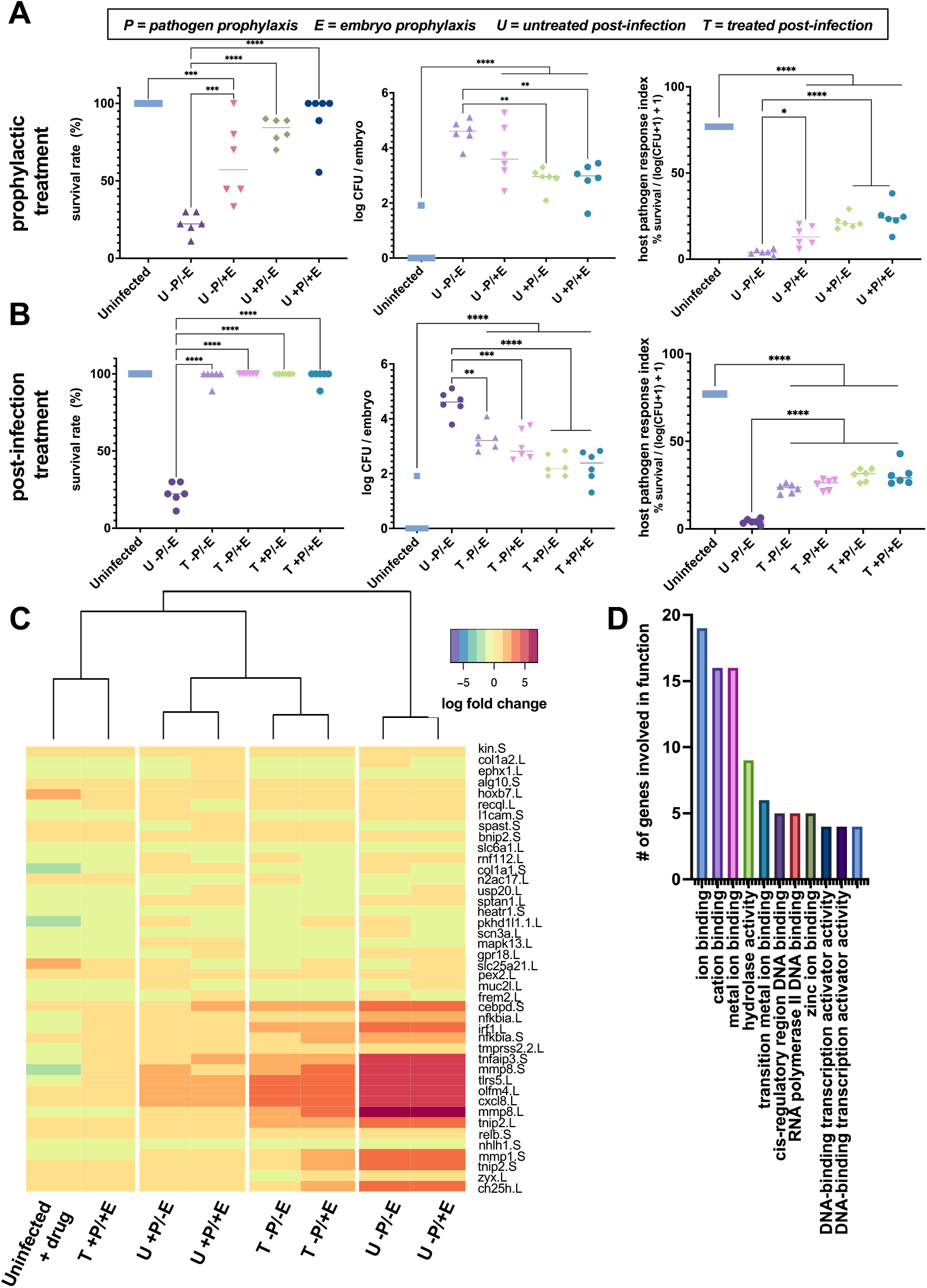
1,4-DPCA treatment mimics the active infection tolerance in the host. **(A)** In groups that received prophylactic treatment, only those in which the pathogens are pretreated (P+) survive at similar rates to controls at 24h post-infection. Embryos treated with prophylactic 1,4-DPCA (U -P/+E) exhibit greater survival than untreated infection (U -P/-E), but lower survival than uninfected controls. Pathogen load decreases with prophylactic treatment of pathogens (**p=0.002), but the load remains higher than that for uninfected controls. The HPRI score is higher for all treatment conditions than untreated infection, including the prophylactic treatment of embryos (*p=0.02), but remains lower than the uninfected controls (***p<0.001; ****p<0.0001; each data point represents n=10 embryos). Statistical analysis was performed using separate one-way ANOVAs for each metric, with post-hoc tests comparing the uninfected controls and untreated infection to each treatment group. **(B)** For groups treated with 1,4-DPCA after the onset of infection (T), survival rates do not differ from uninfected controls at 24h post-infection. Pathogen load decreases in all post-infection treatment groups relative to untreated infection, however, the prophylactically treated and post-infection treated embryos maintain pathogen burdens higher than the uninfected controls. The HPRI score is higher for all treatment conditions than untreated infection, but remains lower than the uninfected controls (**p=0.002; ***p<0.001; ****p<0.0001; each data point represents n=10 embryos). Statistical analysis was performed using separate one-way ANOVAs for each metric, with post-hoc tests comparing the uninfected controls and untreated infection to each treatment group. **(C)** Heatmap comparing the expression of 42 genes that undergo significant changes in gene expression relative to uninfected controls at 24h post-infection (padj<0.001). The prophylactically and post-infection treated group (T +P/+E) clusters with the uninfected control group. Prophylactic treatment of pathogens (+P) and the post-treatment (T) paradigms produce host responses that closely cluster. Embryo prophylaxis alone (U -P/+E) produces a host response that is similar to lack of treatment (U -P/-E). **(D)** Differentially expressed genes are involved in GO pathways related to metal ion binding, hydrolase activity, RNA polymerase binding, and transcription factor activities.

We further investigated how induction of HIF pathways by 1,4-DPCA induces a tolerant state. Transcriptional analysis was carried out in embryos exposed to the prophylaxis and/or post-infection treatment regimens. Hierarchical clustering of differentially expressed genes for each group compared to uninfected controls revealed that prophylactically treating *only* the embryos produces a state similar to that of infected embryos that are not treated with 1,4-DPCA (**Figure 6D**), which is consistent with our finding that this treatment regimen was less effective at inducing tolerance. Similar to earlier assessment of embryos sensitive to infection (**Figure 2A**), prophylactic treatment of embryos only and untreated infection produce marked upregulation of the inflammatory markers CXCL8, TLR5, CEBPD, TNFAIP3, and IRF1 as well as MMPs 1 and 8. Treatment of embryos with 1,4-DPCA after infection produced similar transcriptomic responses that cluster together whether or not the embryos were also exposed to the drug prior to infection (**Figure 6D**). The level of inflammatory markers and matrix metalloproteinases (MMPs) decreased in both these treatment conditions relative to untreated or prophylactically treated embryos. Prophylactic treatment of the pathogen or both the pathogen and the embryo further decreased expression of the inflammatory and MMP genes, and these genes were restored to uninfected control levels by both treating the embryo and pathogen prophylactically and then continuing treatment after initiation of infection (**Figure 6D**). Importantly, the genes that undergo the greatest changes with 1,4-DPCA treatment were involved in cation and metal ion binding (**Figure 6E**), and they are genes that we found within the 20-gene signature for the tolerant state (**Figure 3B**). Other notable changes involved changes in expression of genes in the zinc ion binding pathway, including the zinc-dependent endopeptidases MMP1 and MMP8, as well as the zinc finger protein TNFAIP3. Overall, we show that treatment with 1,4-DPCA induces a host response that closely mimics an active tolerance response when faced with an otherwise lethal pathogen.

## 3. Discussion

In this study, we demonstrated that clinically obtained bacterial pathogens are able to colonize and infect *Xenopus* embryos and stimulate a range of disease responses from passive tolerance to active tolerance or high susceptibility. In infection tolerant *Xenopus* embryos, survival was greater than 90% compared to 0% for susceptible *Xenopus* at 52 hours post-inoculation. Importantly although embryos survived exposure to infection, their pathogen burden remained elevated confirming the induction of a tolerant state rather than an increase in pathogen killing or clearance. To capture this complex host-pathogen state associated with tolerance, we developed the HPRI score, which simultaneously accounts for host survival and pathogen levels in a single score and delineates susceptible, tolerant, and resistant host organisms.

Assessment of gene expression using computational network analysis revealed additional infection tolerance sub-states, with tolerant *Xenopus* embryos clustering into three separate groups at 28 hours post-inoculation: susceptible, actively tolerant, and passively tolerant. Minimal changes in gene expression were observed in the passive tolerance state compared to the activation of many genes within widespread networks in susceptible *Xenopus* embryos. In contrast, we observed more confined gene changes and smaller subnetworks in the active tolerance state, with similar gene expression motifs being observed in tolerant embryos infected with different pathogens. This is similar to the subnetworks observed in the tolerant mouse nasopharynx compared to the very widespread *S. pneumoniae* infection sensitivity in mouse blood^[6]^ and the tightly controlled gene expression observed in mice tolerant to the Ebola virus.^[34]^ Of note, larger changes in *Xenopus* gene expression are seen with infection by gram negative bacteria compared to the gram positive pathogens, so these larger changes may result from LPS or other virulence factors, similar to the large differential gene networks with primates exposed to LPS.^[7]^

Assessment of active tolerance-specific genes and pathways identified changes in ion binding and transport, especially cations and metals, as well as MMPs involved in extracellular matrix remodeling. We also found that the NGB gene that encodes neuroglobin, which is involved in increasing oxygen availability and providing protection under hypoxic and ischemic conditions, ^[30]^ is contained within the tolerance signature. Prior characterization of the transcriptional stress and damage response network associated with systemic infection has similarly implicated iron/heme redox reactions and the hypoxia transcription factor HIF-1α,^[32]^ suggesting their importance in conferring tissue damage control and establishing tolerance to infection.

Broadening our transcriptomic analysis to examine mouse and primate infection models identified overlaps and divergence between mature mammalian responses and those in developing *Xenopus*. Notable overlaps between the mouse nasopharynx and *Xenopus* were observed in their respective tolerance networks with respect to activation of inhibitors within the NF-κB pathway, including IKBKE, which encodes the inhibitor of NF-κB kinase subunit epsilon that regulates inflammatory responses to infection.^[35,36]^ IKBKE also plays an important role in energy balance regulation by sustaining a state of chronic, low-grade inflammation in obesity, which leads to a negative impact on insulin sensitivity.^[21]^ In addition, genes involved in metal ion transport were modulated in tolerance states across *Xenopus*, mice, and primates. These genes included CCND1, a core node previously identified in transcriptional stress and damage responses network^[32]^ and SLC11A1, which can deplete iron and manganese by pumping these metals away from bacteria trapped in the phagosome within macrophages.^[8]^

The overlaps in infection tolerance states across *Xenopus* and mammals combined with the close homology between the human and *Xenopus* genomes^[16,37,38]^ suggests that *Xenopus* could be useful for discovery of therapeutics that induce infection tolerance. *Xenopus* embryos have been previously shown to be a useful model for drug screening^[16,39,40]^ as they enable rapid evaluation of drug efficacy due to their small size, extrauterine development, skin permeability to small molecules, and relative high-throughput with thousands of eggs from a single female.

Based on the pathways that we found define the tolerance state in *Xenopus* and analysis of known mechanisms of tolerance, we screened drugs in *Xenopus* embryos infected with *A. hydrophila* bacteria, which induce high mortality levels in the absence of treatment. Both the metal ion chelator, DFOA, and the HIF agonist, 1,4-DPCA, were found to shift *Xenopus* to a tolerant state as indicated by an increase in survival despite not clearing the bacterial pathogen. Although prophylaxis with 1,4-DPCA prior to bacterial infection suggested there is some effect on bacteria, prophylactic treatment of embryos alone improves survival and post-infection treatment is the most effective, clarifying the critical role of the host response to these drugs.

Hypoxia and metal ion binding pathways are interconnected^[8]^ and therefore targeting of either can induce tolerance to infection. Both metal ions and functional prolyl 4-hydrolase enzymes are required to inhibit the production of HIF, and metal ion chelators and scavengers compete with collagen prolyl hydroxylase (CPH) for free metals. Thus, these treatments result in inhibition of CPH-mediated hydroxylation of HIF-1α, which prevents HIF-1α degradation (**Figure S5**). Preventing this HIF degradation may reduce tissue damage and increase disease tolerance to infection by metabolic adaptation in glucose, metabolites, and metals like iron that modulate macrophage polarization.^[1,41]^

In addition to its role in HIF pathways, scavenging metal ions can lead to nutritional immunity by starving pathogens of essential nutrients for biological processes.^[1,8]^ Bacteria require metal ions as cofactors or structural elements for enzymes, metabolic function, and virulence factor expression,^[8]^ which may explain the success of metal scavenging approaches for inducing a tolerance state (**Figure S5**). Furthermore, connections exist between circadian, metal, and hypoxia pathways that may account for their shared ability to produce a tolerant state. We observed some evidence of these connections in *Xenopus*, with metal ion binding, an oxygen transporter, and the circadian gene HNF4A all being included within the tolerance gene signature.

HNF4A represses CLOCK-ARNTL/BMAL1 transcriptional activity and is essential for circadian rhythm maintenance and period regulation in liver and colon cells.^[24]^ The connections between circadian rhythm and metal ion levels, such as iron, can be explained by the central role of heme, an iron-containing porphyrin that reversibly binds to nuclear receptor Rev-erbα, a critical negative component of the circadian core clock.^[42,43]^ Heme also acts as prosthetic group for enzymes involved in oxidative metabolism and transcription factors that regulate circadian rhythmicity.^[42]^ In addition, hypoxia directly impacts circadian gene expression through the production of acid that suppresses the circadian clock through diminished translation of clock components.^[33]^ The coordination between circadian cycles and the immune system may be especially important for promoting rhythms in immunity that anticipate exogenous microbial exposure through oscillations in epithelial cell STAT3 expression, as demonstrated in the rodent in response to intestinal microbiota.^[44]^ Because we demonstrate that host tolerance can be induced through at least three pathways, there might be an opportunity for development a new class of tolerance-inducing drugs that could minimize tissue damage in the host during infection in the future.

We anticipate that host tolerance will be a valuable strategy to reduce the morbidity and even mortality associated with acute infection, such as bacterial septicemia, when it is not possible to achieve sterile immunity. For example, recent work has shown that immunity to severe life-threatening infection with malaria is underpinned by acquired mechanisms of disease tolerance^[45,46]^, and this new understanding suggests that a future focus on tolerance-inducing treatments is warranted. The use of a broad-spectrum host tolerance-inducing drug may be especially useful before a vaccine is available for a novel infectious pathogen or when bacteria develop resistance to existing antibiotics or antivirals. Although the present study focused on a single treatment of tolerance-inducing compounds, we anticipate that these drugs might synergize with antibiotic and antiviral therapies when used in combination by both inducing tolerance and reducing the burden of pathogens, which would decrease the likelihood of spreading to others while also further reducing tissue damage. However, infection tolerance could result in organisms never fully clearing the pathogen and persistence of low-grade infection. From a public health perspective, a tolerant individual could also continue to spread the infection and endanger non-tolerant individuals around them.^[47]^ Therefore, careful consideration of public health outcomes must be considered before translation of tolerance drugs into human patients.

A unique aspect of Xenopus embryos is the lack of adaptive immunity development, enabling the study of innate immune responses in the presence of a complex organ system interaction without the overlapping adaptive response. Although immature *Xenopus* elicit strong immune responses,^[17]^ their responses are different and therefore any insights from the *Xenopus* embryo model will need to be validated in more adult-like infection models. Nevertheless, the success of this integrated experimental-bioinformatics screening approach used here demonstrates the value of the *Xenopus* infection model for gaining insight into mechanisms of host tolerance to infection and for discovery of broad-spectrum tolerance-inducing drugs. Using transcriptomic signatures and the *Xenopus* embryo infection screen, we identified that the hypoxia pathway involving prolyl-4 hydroxylase and HIF-1α play key roles in modulating infection tolerance, and showed that metal ion chelators and a prolyl-4 hydroxylase inhibitor can act as potential tolerance inducers that mimic an active tolerance response to infection. Thus, this combined experimental and computational platform may help to accelerate the development of a new class of tolerance-inducing therapeutics that could complement antibiotic therapies in the fight against life-threatening infectious diseases, such as bacterial septicemia.

## 4. Methods

### Xenopus husbandry

All experimental procedures involving *Xenopus* embryos were approved by the Institutional Animal Care and Use Committees (IACUC) and Tufts University Department of Laboratory Animal Medicine under protocol M2014-79 and by the Office of the IACUC at Harvard Medical School under protocol IS00000658-3. *Xenopus laevis* embryos were fertilized *in vitro* according to standard protocols in 0.1X Marc’s Modified Ringer’s solution (MMR; 10 mM Na^+^, 0.2 mM K^+^, 10.5 mM Cl^−^, 0.2 mM Ca^2+^, pH 7.8) and housed at 18°C. Since exclusively sterile filtered solutions, including MMR culture medium, were used to avoid confounding effects of seasonally-varying environmental bacteria, we dosed embryos overnight with 2.64 μl KoiZyme probiotic solution (Koi Care Kennel, Las Vegas, NV) per 1L of 0.1% MMR to avoid dysbiosis (**Figure S6**). Sterile solutions were used for all studies. Embryos were staged according to Nieuwkoop and Faber.^[48]^

### Microinjection of clinically-obtained pathogen strains

Bacteria (**Table 1**) were streaked out from frozen glycerol stock of target bacterial strain on sheep blood agar media plates overnight at 37°C. A single colony was selected from the streak and grown in LB medium at 37°C. Exponentially growing bacteria were pelleted and resuspended in sterile saline/dextrose with 15% glycerol at a concentration of 1×10^9^ CFU/ml. Faber-Nieuwkoop stage 13/14 embryos were injected with fresh bacterial suspensions using borosilicate glass needles calibrated for a bubble pressure of 25–30 kDa and 0.4 sec pulses to deliver 10^3^–10^4^ CFU of bacteria to each embryo (**Figure S1**). Bacterial injections were performed with embryos submerged in 3% Ficoll prepared in 0.1X MMR. Even though bacteria were grown and concentrated following a constant methodology and embryos were collected and grown under the same conditions, variation in survival rates was observed between experiments. This is likely due to differences in bacterial growth *in situ*, which can be affected by differences in the genetic background of each individual from different egg clutches. For these reasons, every individual comparison within an experiment was conducted using eggs from a single fertilization and a single suspension of bacteria and replicate studies were conducted with completely independent batches of embryos.

**Table 1.** Pathogen Sources.

| Pathogen | Source |
| --- | --- |
| *S. aureus* 3518 | Crimson |
| *P. aeruginosa* 41504 | Crimson |
| *A. baumannii* | Crimson |
| *A. hydrophila* | Crimson |
| *S. pneumoniae* | Crimson |
| *K. pneumoniae* | Boston Children's Hospital |

**Table 2.**
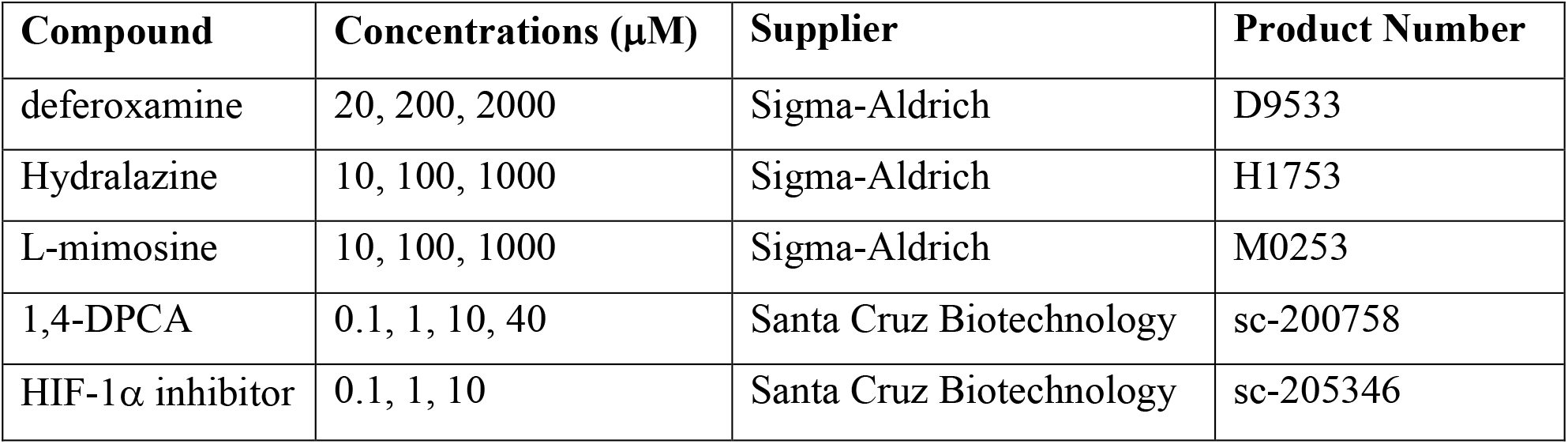
Key Compounds & Concentrations Tested.

### Quantification of host infection tolerance

Infection tolerance state was determined by a combination of the survival rate and pathogen burden in a group of embryos. At selected time points after infection, embryo survival was assessed by microscopic evaluation. At the same time points, embryo lysate was spiral plated onto selective media plates by the Eddy Jett 2 Spiral Plater, cultured, and counted using the Flash & Go Automated Colony Counter to determine the concentration of viable pathogen. The survival and amount of viable pathogen can then be combined into a single score that reflects the host infection tolerance state, the Host Pathogen Response Index:

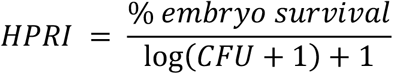

### Drug screening

Embryos were exposed to ion scavenging and hypoxia-inducing compounds from Faber-Nieuwkoop stage 13/14, including DFOA (Sigma-Aldrich; St. Louis, MO), hydralazine (Sigma-Aldrich), L-mimosine (Sigma-Aldrich), and 1,4-DPCA (Santa Cruz Biotechnology; Dallas, TX) (**Figure S1**). Compounds were dissolved in DMSO and further diluted in 0.1X MMR to a range of micromolar concentrations (**Table 2**). Embryos infected with *A. Hydrophila* were dosed with drugs in 12-well plates (n=10 embryos/well) to assess therapeutic potential. Embryo survival, levels of viable pathogen, and HPRI were assessed for each treatment at 24, 48, and 120 hours post-infection. To test the primary mechanism of action for 1,4-DPCA, embryos were co-treated with 1,4-DPCA and a HIF-1α inhibitor (Santa Cruz Biotechnology). 1,4-DPCA treatment was also tested across the circadian cycle by flipping light cycles from the normal 12/12 light/dark cycle to a 12/12 dark/light cycle using isolated light boxes within the culture incubators. Following initial drug screening and mechanism studies, prophylactic and post-infection treatment were compared for 40μM 1,4-DPCA treatment at 24h post-infection. Statistics were calculated and plots generated using Prism Version 9.1.2 (GraphPad; San Diego, CA).

### Transcriptomics

To determine the effects of bacterial infection and treatments on expression profiles in embryos, microarray analysis was performed at the conclusion of two experiments (**Figure S1**). Separate microarray analyses were used to assess: 1) The effect of clinically-obtained pathogen strains (**Table 1**) on embryos at 4h and 28h post-infection and 2) The therapeutic effect of prophylactic and post-infection treatment with 1,4-DPCA on embryos infected with *A. hydrophila* at 24h post-infection. For each experiment, RNA from each *Xenopus* embryo (n=2-3/condition) was separately extracted and purified using the RNeasy Micro Kit (Qiagen; Venlo, Netherlands) and microarray measurements were performed using the GeneChip *Xenopus laevis* Genome 2.0 Array (Affymetrix; Santa Clara, CA) at the Advanced Biomedical Laboratories (Cinnaminson, NJ). Microarray data were extracted from CEL files, Robust Multi-array Average (RMA)^[49,50]^ normalized in Matlab (Mathworks; Natick, MA), and expression data was log2-transformed. The limma package^[51]^ in R was used to determine differentially expressed genes after infection and treatment relative to uninfected controls using the Benjamini-Hochberg false discovery rate (FDR).^[52]^ Heatmaps displaying genes that undergo the greatest differential expression were produced using the gplots package.^[53]^ To identify tolerant-specific genes, we filtered for genes that were differentially expressed (FDR<0.05) in the active tolerance state (*K. pneumoniae* and *A. baumannii* infections) and removed any genes that were differentially expressed in the sensitive state (*P. aeuruginosa* and *A. hydrophila* infections).

### Pathway Analyses

The infection tolerance expression signature was assessed using gene ontology (GO) enrichment tools to highlight pathways relevant to infection tolerance. Tolerance-specific *Xenopus* gene names were first converted to human orthologs based on the HGNC Comparison of Orthology Predictions (HUGO Gene Nomenclature Committee at the European Bioinformatics Institute; https://www.genenames.org/). The PANTHER Classification System^[26]^ (http://pantherdb.org/) was used to perform a functional classification analysis based on GO^[27,28]^ biological processes, molecular function, and Reactome^[29]^ (https://reactome.org/) pathways amongst genes of the tolerance signature for *Homo sapiens*. Pathways were identified using the PANTHER Overrepresentation Test and sorted based on the number of genes identified along each pathway.

### Gene network analyses

Gene networks for each bacterial infection were analyzed and visualized in Matlab (Mathworks; Natick, MA) to identify subnetworks and motifs of interacting genes that are activated in infection tolerance and sensitive states. The overall gene network was built using known molecular interactions from the KEGG (https://www.genome.jp/kegg/) and TRRUST databases (https://www.grnpedia.org/trrust/).^[18–20]^ The active subnetworks and motifs for each infection and treatment condition were identified by searching the overall network for genes found to be significantly differentially expressed by limma analyses. Activated subnetworks and motifs include genes that are interconnected with at least one other gene and disconnected gene nodes were removed from each network. For each network, the importance of each node was determined by calculating its degree centrality, which counts the number of edges connecting to each node. The same methods were applied to previously published and processed gene expression data from *S. pneumoniae*-infected mice^[6]^ and LPS-exposed primates^[7]^.

## Data availability

Microarray data reported in this paper will be deposited in the GEO database (https://www.ncbi.nlm.nih.gov/geo/) upon publication.

## Supporting Information

Additional data is available in the Supplementary Materials.

## Acknowledgements

The authors acknowledge funding from DARPA (W911NF-16-C-0050) and thank E. Switzer and E. Lederer for help with *Xenopus* embryo fertilization and husbandry and B. Melakeberhan for assay protocol support. Biorender (https://biorender.com/) was used for the making of all schematic figures.

