## Supplemental Figures for "Enhancers of host immune tolerance to bacterial infection discovered using linked computational and experimental approaches"

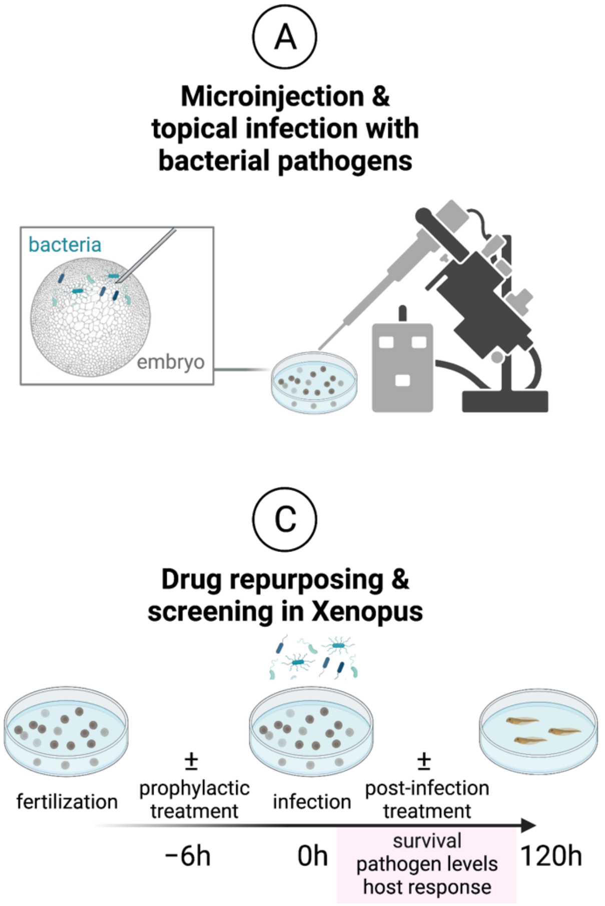


**Figure S1. Using a combination of bioinformatics and experimental approaches with Xenopus embryos enables rapid discovery of relevant pathways, repurposing of candidate drugs, and execution of efficacy and toxicity screening to identify compounds that induce broad tolerance to bacterial infection.** Embryos were first injected or topically infected with bacterial pathogens (**A**). Gene expression was then assessed in infected embryos by transcriptomics, pathway enrichment, and gene network analyses (**B**). Based on bioinformatics outcomes, drugs were selected for repurposing to target relevant pathways and screened in infected Xenopus embryos (**C**). For compounds that produced infection tolerance states, targets were verified using a combination of transcriptomics and inhibitors that blocked relevant pathways (**D**).

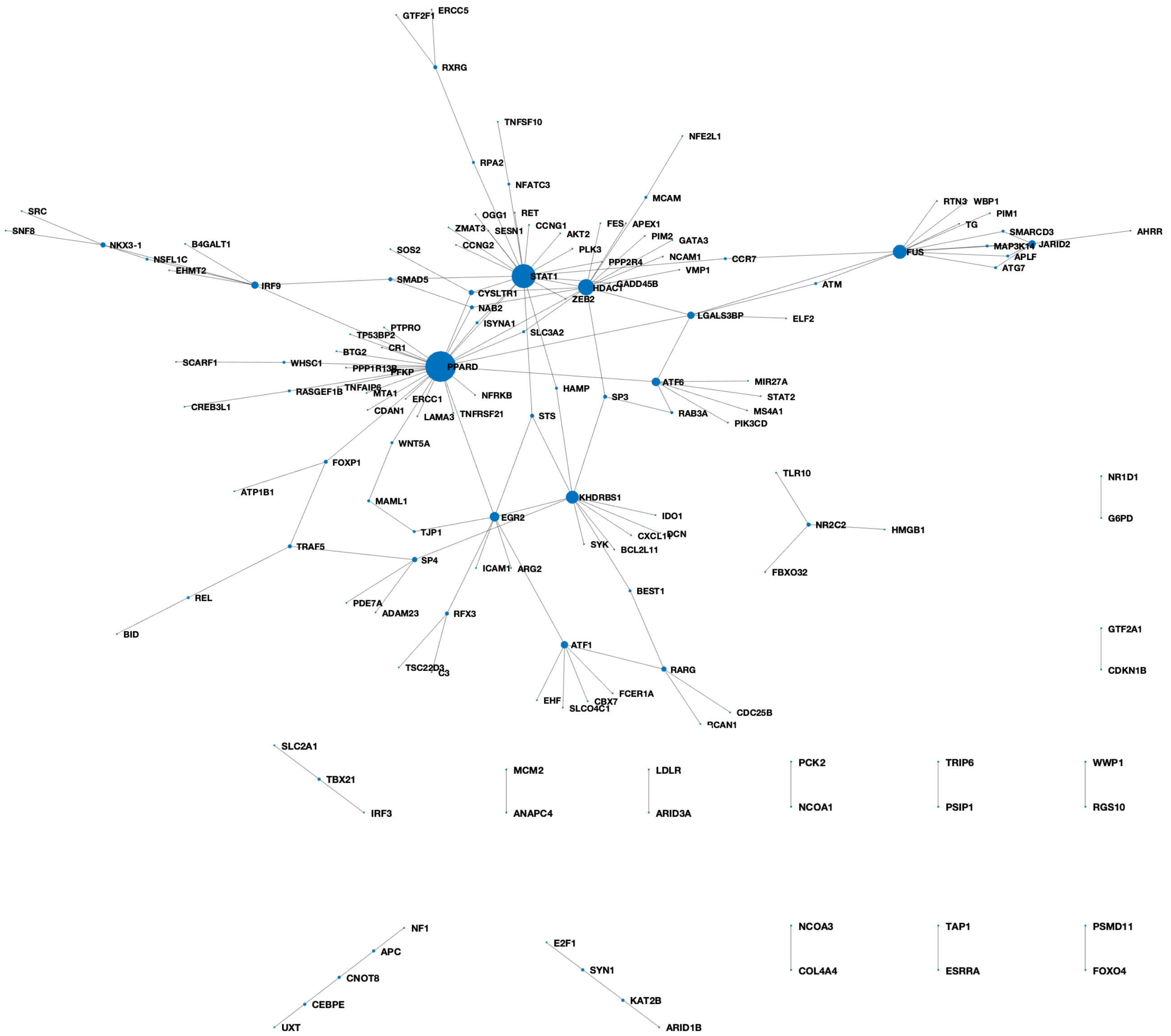


**Figure S2.** Tolerance-specific gene networks in primates are comprised of genes involved in interferon alpha and gamma regulation, T cell activity, response to metal ions, and cellular hypoxia.

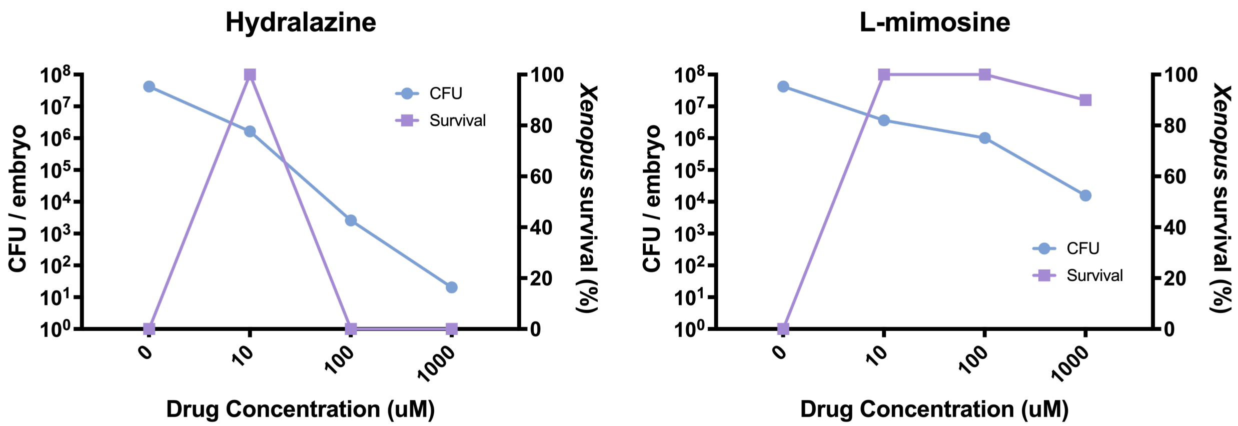


**Figure S3.** The metal scavengers hydralazine and L-mimosine improve *Xenopus* survival but minimally impact pathogen load at non-toxic doses, thereby inducing a tolerant state. Hydralazine substantially impacts pathogen levels only at high drug concentrations that are toxic for *Xenopus* embryos.

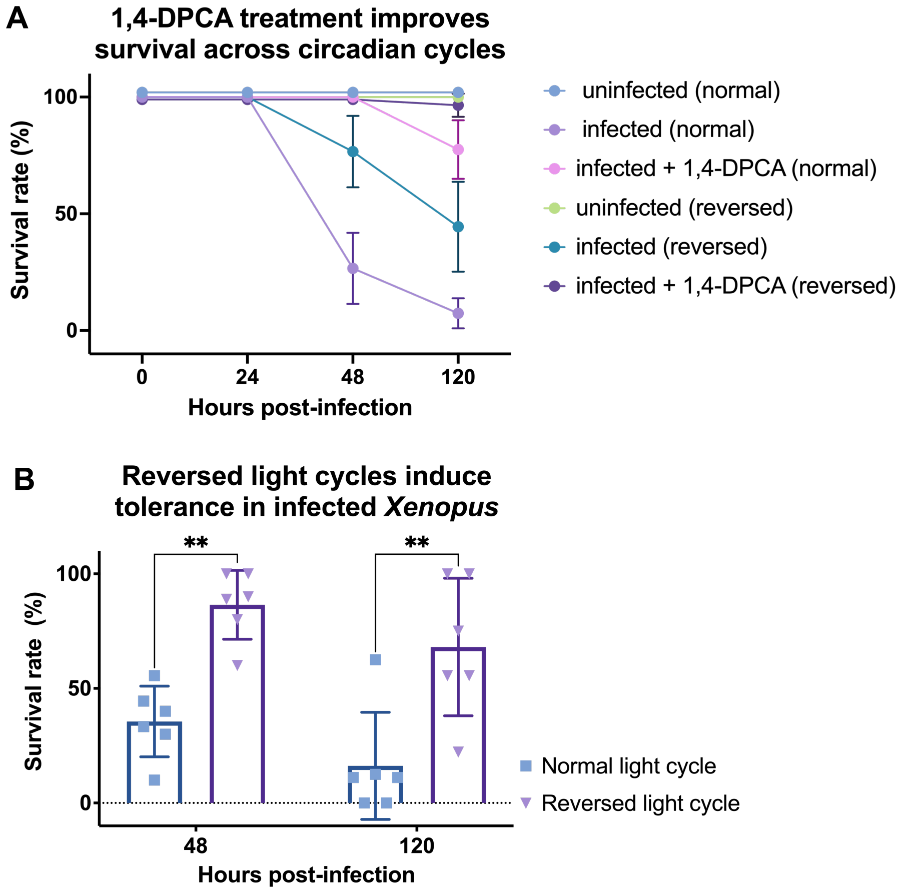


**Figure S4. Light cycle perturbation induces tolerance.** (**A**) 1,4-DPCA is capable of inducing tolerance regardless of light cycle, with no differences detected between the normal 12/12 light-dark cycle and the reversed 12/12 dark-light cycle. (**B**) In infected, untreated *Xenopus*, flipping the light cycle improved tolerance to *A. hydrophila* infection, which is not observed in *Xenopus* exposed to a normal light cycle (**p<0.01). Statistical analysis was performed using a repeated measures ANOVA and Sidak's test was used to correct for multiple comparisons.

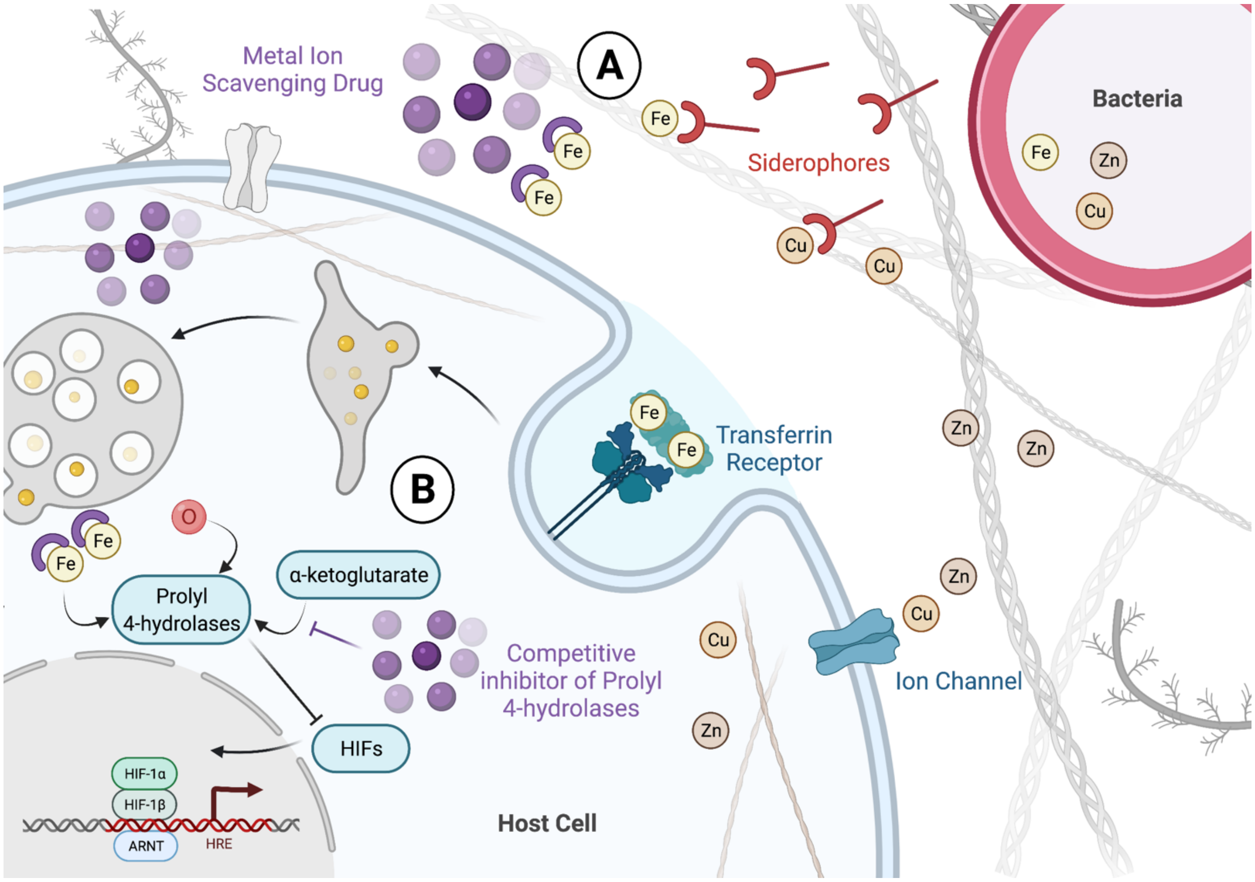


**Figure S5. Artificial induction of infection tolerance by modulation of cellular hypoxia response.** DFOA, hydralazine, and L-mimosine function as metal ion scavengers **(A)**. Scavenging of metal ions prevents the uptake of ions by bacteria via sidephores, a molecule that binds and transports iron in microorganisms. Metal ions are necessary for enzyme function, normal metabolic activities, and virulence factor expression in bacterial pathogens. Scavenging of metal ions, particularly iron, also prevents the function of prolyl 4-hydrolases in degrading HIF proteins required for production of hypoxia responsive elements (HREs). **(B)** Competitive inhibition of prolyl 4-hydrolaseswith via 1,4-DPCA, an a-ketoglutarate mimic, also modulates the same pathway by increasing HIF protein production and the downstream HREs.

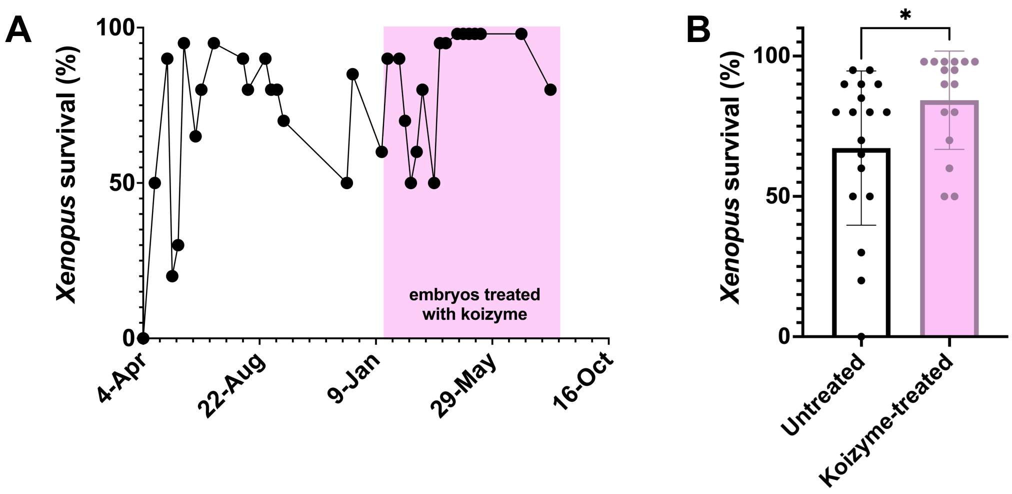


**Figure S6. Koizyme improves survival in *Xenopus* embryos by preventing dysbiosis.** (**A**) The overnight viability of embryos after fertilization for experiments over a 1.25-year time span. The period of Koizyme use is highlighted in pink. (**B**) The addition of Koizyme to *Xenopus* media overnight after fertilization improves embryo survival over untreated embryos (*p=0.04). Statistical analysis was performed using a two-tailed Student's *t* test.
